# Cooperative molecular interaction networks govern PARP1 inhibitor selectivity and binding affinity

**DOI:** 10.1101/2025.10.13.681983

**Authors:** Alejandro Feito, Natàlia DeMoya-Valenzuela, Cristian Privat, Andrés R. Tejedor, Lucía Paniagua-Herranz, Adiran Garaizar, Alberto Ocana, Jorge R. Espinosa

## Abstract

Selective inhibition of PARP1 represents a promising strategy to improve the therapeutic index of PARP inhibitors, a class of anticancer agents that exploit defects in DNA repair pathways. While PARP inhibitors have shown remarkable clinical benefit, particularly in BRCA-mutated tumors, the lack of discrimination between PARP1 and its close homolog PARP2, often leads to hematological toxicity and limits treatment efficacy. Thus, achieving molecular selectivity for PARP1 remains a central challenge in the rational design of safer and more potent inhibitors. To explore the molecular determinants of ligand selectivity, we focus on four clinically relevant PARP inhibitors—two PARP1-selective (saruparib and NMS-P118) and two non-selective (veliparib and olaparib) inhibitors—and perform atomistic potential-of-mean-force calculations of the PARP1 catalytic binding domain in the presence of these molecules. Our simulations near-quantitatively capture the experimental relative binding preferences, demonstrating that our approach reliably reflects selectivity patterns. Based on these findings, we analyze protein–ligand contact frequencies to identify the stabilizing interaction network and contact connectivity inducing protein selectivity. The most frequent protein–inhibitor contacts are primarily mediated by tyrosine triads and electrostatic interactions, showing a cooperative complex network of intermolecular contacts which strongly relies on protein multivalency. To dissect the decisive role of individual residues across the binding site, we also perform targeted mutagenesis of the PARP1 catalytic pocket in complex with saruparib, replacing several active-site amino acids by glycines. Progressively increasing the number of mutations markedly reduces binding stability, with distinct residue combinations exerting two primary effects: destabilization of the final bound state and the emergence of energetic barriers along the ligand association pathway. Together, our results provide a coherent mechanistic framework for understanding PARP1 selectivity and informs the rational design of next-generation inhibitors with improved efficacy and safety.

## I. INTRODUCTION

Poly(ADP-ribose) polymerase 1 (PARP1) and 2 (PARP2) are nuclear enzymes that play essential roles in the detection and repair of DNA damage^1,2^, particularly single-strand breaks^3^. Acting as central guardians of genomic integrity, these enzymes orchestrate coordinated responses to DNA lesions that ensure efficient repair^2^, preserve chromatin organization^2^ and maintain cellular viability^4^. Upon sensing DNA damage, PARP1—responsible for the majority of damage-induced PARylation—and the related enzyme PARP2 catalyze the polymerization of poly(ADP-ribose) chains, thereby recruiting nuclear factors that promote chromatin remodeling^5–7^ and the assembly of DNA repair complexes^8^. Beyond their canonical roles in DNA repair, PARP1 and PARP2 contribute to transcriptional regulation^9^, replication fork stability^10^, and cellular responses to oxidative stress^11^, underscoring their broad involvement in nuclear homeostasis. Defects in these pathways—such as those associated with *BRCA1/2* mutations^12,13^—are tightly linked to oncogenesis^14–18^. Therefore, the central role of PARP1, and to a lesser extent PARP2, in DNA damage sensing and repair makes them compelling therapeutic targets, particularly in tumors dependent on compensatory repair mechanisms for survival^19,20^.

Targeting PARP1 with small-molecule inhibitors has become an effective therapeutic strategy^21–24^, exploiting synthetic lethality in cancers with homologous recombination (HR) repair deficiency^25,26^. Clinically approved PARP inhibitors—including olaparib^27–29^, rucaparib^30,31^, niraparib^32,33^, and talazoparib^34,35^—inhibit both PARP1 and PARP2. This lack of selectivity contributes to a narrow therapeutic window and characteristic hematologic toxicities largely attributed to PARP2 inhibition^14,29^ . Next-generation inhibitors such as saruparib^37^ or NMS-P118^38^ were designed to achieve greater PARP1 selectivity, thus minimizing adverse hematologic effects^39^ and emphasizing the clinical importance of selectively targeting PARP1^18^. Developing highly specific PARP inhibitors not only promises improved tolerability but also enables rational combination therapies with additional DNA-damaging agents under a safer therapeutic profile^40^. Both saruparib and NMS-P118 were conceived as PARP1-selective inhibitors, but their clinical outcomes differ: NMS-P118 displayed approximately 150-fold selectivity over PARP2 and remained preclinical^38^, while saruparib exhibited up to 500-fold selectivity^41^, enhanced hematologic safety, and has advanced to clinical trials^42^, showing durable responses in HR-deficient tumors^18^.

Understanding the molecular basis of inhibitor selectivity requires mechanistic knowledge of how these ligands interact with specific amino acid residues within the catalytic binding site, as well as with accessory factors such as HPF1^43,44^, which modulate enzymatic activity^18,45–47^. Although high-resolution structural data provide essential configurations of these interactions, they often fail to capture the conformational dynamics and solvent effects which are critical for predicting binding affinity and specificity^48,49^. Atomistic molecular dynamics (MD) simulations^50–54^ can successfully address these limitations by describing inhibitor binding at the atomistic level in an explicitly solvated medium and within a dynamically evolving environment under physiological conditions. By characterizing the conformational flexibility of the active site, identifying key protein-ligand contacts, and quantifying binding energy landscapes^55^, MD simulations bridge static structural information with the dynamic behavior of biomolecular systems. Unlike crystallographic models, these simulations incorporate solvent, ions, and conformational ensembles, producing a realistic description of ligand–protein recognition. Such computational framework enables the dissection of the energetic and structural determinants underlying potency and selectivity, allowing systematic testing of hypotheses which are often inaccessible experimentally. In this way, MD simulations not only guide rational drug design and optimization of existing inhibitors, but can also accelerate the discovery of new compounds with improved efficacy and safety.

In this study, we perform atomistic MD simulations of PARP1 and PARP2 catalytic binding domains in complex with a representative set of inhibitors, including PARP1-selective compounds (NMS-P118^38^ and saruparib^37^) and non-selective inhibitors (olaparib^27^ and veliparib^27,56^). The binding free energy is evaluated via potential-of-mean-force (PMF) calculations, while residue-level protein-ligand contacts within the catalytic binding site are analyzed and compared with available experimental data. To connect bound-state interactions with ligand binding strength, we further perform targeted *in silico* mutagenesis by replacing selected activesite amino acids by glycines, with particular emphasis on the interactions driving selectivity in saruparib. Together, our results integrate structural, energetic, and residue-level analyses to elucidate the molecular determinants of PARP1 selectivity, providing a mechanistic framework for the rational design of more potent and specific PARP1 inhibitors.

## RESULTS AND DISCUSSION

### Potential of Mean Force Calculations to Quantify PARP–Ligand Interactions: Insights into the Binding Mechanisms and Ligand Selectivity

To quantify the strength of ligand-protein binding interactions and to characterize the energetic pathway underlying ligand association within the active site, we perform PMF calculations using Umbrella Sampling simulations^57–60^ and the all-atom a99SB-*disp* force field with the TIP4P-*disp* water model^61,62^ under physiological conditions (310 K, 150 mM NaCl; see Sections SI and SII of the Supplementary Material (SM) for further methodological details). The force field parameters and topology for all ligands are derived using the OpenFF toolkit^63^, ensuring full compatibility with the all-atom parameters of the protein sequence using the a99SB-*disp* force field (see Section SI of the SM for further computational details). Unlike static structural analysis, this approach allows us to explicitly probe the free energy landscape governing the association and dissociation pathways of the different inhibitors. PMF simulations provide a quantitative description of the free energy profile along a defined reaction coordinate, chosen here as the center-of-mass (COM) distance between the inhibitor and the catalytic binding pocket. This enables direct monitoring of how the interaction energy evolves as the ligand approaches or moves away from the protein binding site (see Section SI in the SM for simulation details of this method). Our PMF initial configurations are build from crystallographically resolved protein–ligand complexes available in the Protein Data Bank (see Section SII of the SM for PDB codes used in this study), with the exception of the PARP2–saruparib system, for which no experimental structure is available. For this system, the initial bound configuration is generated through molecular docking structural alignment into the corresponding PARP2 pocket^64^. From these initial configurations for each PARP1/PARP2-ligand complex, we generate a series of protein–ligand subsequent configurations separated by incremental COM distances and ensuring overlap along the reaction coordinate, defining different Umbrella Sampling windows. For each window, we perform independent simulations subsequently combined to reconstruct the PMF profile as a function of the protein–ligand COM distance (Figure 1A). The resulting PMF profiles exhibit a pronounced free energy minimum near the crystallographic binding distance, corresponding to the most stable bound configuration. As the ligand is progressively displaced from the binding site, the free energy increases and eventually plateaus near zero, indicating complete dissociation and the absence of effective protein–ligand intermolecular interactions beyond such distance (see Figure 1A for a schematic representation of a ligand-protein PMF profile).

**FIG. 1.**
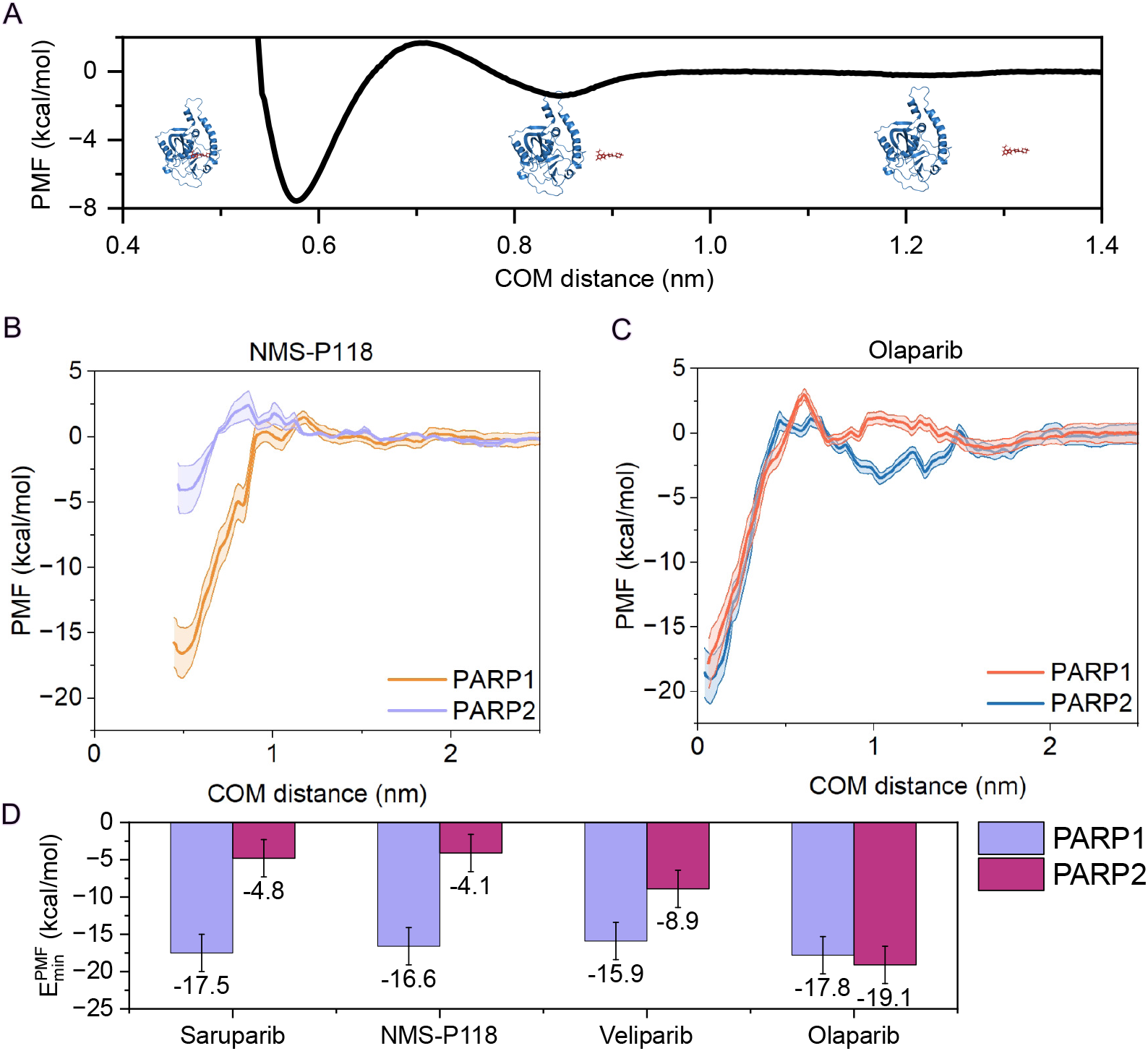
(A) Schematic representation of a PMF profile between the protein and the ligand as a function of the distance of the center of mass (COM). Inset show representative configurations taken at different protein–ligand distances along the reaction coordinate (COM distance). (B–C) Atomistic PMF dissociation profiles between each ligand and PARP1/PARP2 under physiological NaCl concentration (150 mM) and room conditions, in explicit solvent and ions. The COM distance between the protein binding pocket and the ligand is used as the reaction coordinate. Curves are shown for NMS-P118 (B) and olaparib (C), depicting the statistical uncertainty of each PMF profile by a color shadowing corresponding to the standard deviation obtained from trajectory segments (see Section SI of the SM for further details). (D) Summary of the free energy minima obtained through PMF calculations for all ligands in complex with PARP1 (violet) and PARP2 (maroon).

In Figure 1B–C, we present the PMF profiles for NMS-P118 and olaparib, respectively, including the corresponding curves for PARP1 and PARP2. The PMF for NMS-P118-PARP1 (orange curve; Figure 1B) exhibits a deep energy minimum of approximately -16.6 kcal/mol, indicative of high binding affinity. The energy landscape is relatively flat, showing significantly low desolvation energy barriers at intermediate distances (1–2*∼* Å). In contrast, the PMF for PARP2 (violet curve) reveals a markedly weaker binding affinity, with a shallow minimum of only -4.1 kcal/mol. Notably, this profile features a moderate energy barrier of approximately 3 kcal/mol at intermediate distances (0.8 Å) that is absent in PARP1. Such barrier possibly reflects unfavorable desolvation effects and the energetic cost of displacing water molecules solvating NMS-P118 prior to the formation of energetically favorable ligand–protein interactions, thereby hindering the association of NMS-P118 with PARP2. Although the final bound state remains moderately favorable, the combination of a weak minimum for PARP2 and a much higher binding free energy for PARP1 explains the experimentally established selectivity of NMS-P118 for PARP1^38,65^. For olaparib (Figure 1C), the PMF profile for PARP1 (red curve) shows an energy minimum of -17.8 kcal/mol, while PARP2 (blue curve) exhibits a similarly deep minimum of -18.1 kcal/mol. Both curves gradually descend along the reaction coordinate, indicating low to moderate energy barriers and an overall smooth association pathway. Notably, the PARP2 curve displays a subtle local minimum at intermediate distances, suggesting transient interactions that may guide the ligand toward the active site and facilitate binding. The comparable depths of the energy wells in both protein pockets provide a clear mechanistic explanation for the lack of selectivity of olaparib, consistent with experimental measurements^66^. The same calculations are also performed for both saruparib and veliparib (Figure S1 of the SM), and the depth of the global minimum for each PMF profile is reported in Figure 1D.

Our results are broadly consistent with experimental observations: both saruparib and NMS-P118 exhibit a markedly higher binding affinity for PARP1, veliparib shows a modest preference for PARP1^67,68^, while olaparib is generally considered non-selective^68,69^ (Figure 1D). Measurements of the half maximal inhibitory concentration, IC_50_, indicate that both veliparib and olaparib only show limited PARP1 selectivity and, in certain contexts, even a relative preference for PARP2^65,66,68,70–72^.We provide a direct comparison between the binding free energy values (Δ*G*) obtained from our PMF profiles and those experimentally derived from IC_50_ values using the following relation^73^:

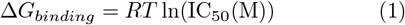

where *R* is the gas constant, and *T* = 300 K. Experimental IC_50_ values were taken as follows: saruparib, 3 nM for PARP1 (closed blue symbol) and 1400 nM for PARP2 (open blue)^41^; NMS-P118, 9 nM for PARP1 (closed red) and 1390 nM for PARP2 (open red)^38^; veliparib, 8.3 nM for PARP1 (closed green) and 11 nM for PARP2 (open green)^71^; and olaparib, 1.1 nM for PARP1 (closed orange) and 0.9 nM for PARP2 (open orange)^74^ and then converted into binding Δ*G* values. In Figure 2, the experimentally derived Δ*G* values are compared with those extracted from the PMF profiles, revealing a qualitative correlation across the data set. We note that the absolute free energies quantitatively differ: experimental values of Δ*G* range approximately from *−*8 to *−*13 kcal mol*^−1^*, whereas the PMF minima span from about *−*20 to *−*5 kcal mol*^−1^*. This discrepancy likely arises from several potential sources: (1) inherent limitations of the employed force field^61^; (2) the positional restraints applied during the PMF calculations which constrain the ligand conformational freedom and may thus overstabilize the binding interaction strength in all systems^59,75^; and (3) the choice of the reaction coordinate for the dissociation pathway along the PMF calculations. All these factors introduce a systematic offset that cannot be precisely quantified. Therefore, the error bars in Figure 2 reflect only the statistical uncertainty of our PMF calculations. Nevertheless, despite these sources of uncertainty and our systematic offset (which would affect all systems equally), our simulations near-quantitatively reproduce the experimentally reported selectivity for these ligands. Overall, the relative binding preferences between PARP1 and PARP2 are consistently captured, confirming that our PMF-based approach reliably recapitulates the experimental selectivity patterns. A similar trend to that shown in Figure 2 can also be observed in Figure S2 of the SM, which compares the differences in Δ*G*_binding_ between PARP1 and PARP2 for both experimentally and computationally-derived free energies.

**FIG. 2.**
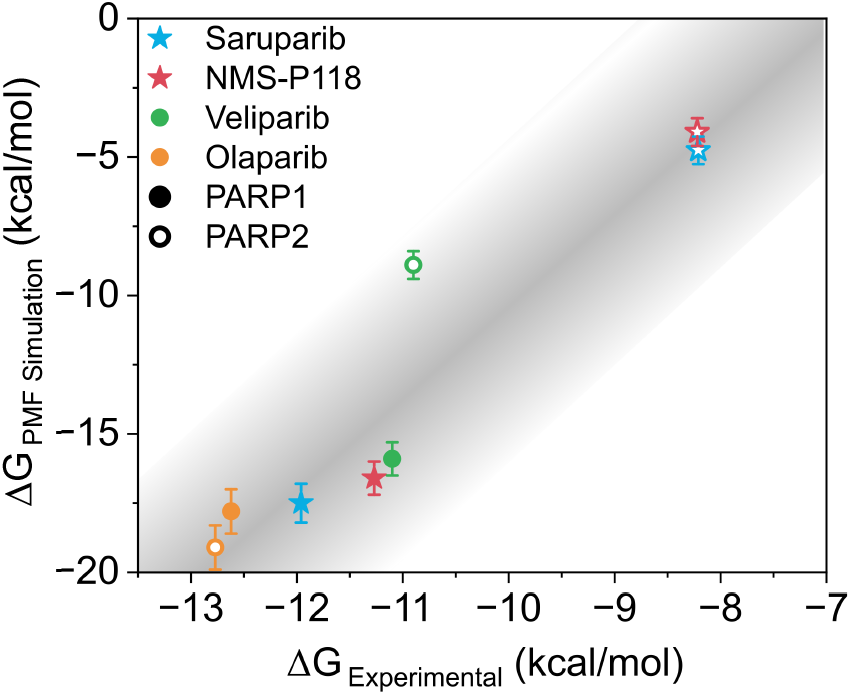
Comparison between experimental (Δ*G*_Experimental_) and simulated (Δ*G*_PMF Simulation_) binding free energies for different inhibitors (Saruparib, NMS-P118, Veliparib, and Olaparib) for PARP1 (closed symbols) and PARP2 (empty symbols). Specific inhibitors of PARP1 are plotted in stars and non-specific inhibitors in circles. A linear shading to the obtained correlation was added as a guide to the eye.

### Residue-level Contact Analysis of PARP Inhibitors

Understanding atomic-level interactions between small-molecule inhibitors and the active site of PARP1 *vs.* PARP2 is essential to elucidate the molecular basis of selectivity. Consequently, we analyze the molecular contact network between the selected ligands and the catalytic site of both enzymes obtained from 1 microsecond trajectories (see Section SIII of the SM for further details). We select two PARP1-specific inhibitors (NMS-P118^38^ and saruparib^37^) and two non-selective inhibitors (olaparib^27^ and veliparib^27,56^) to assess whether the resulting contact profiles can provide insight into ligand selectivity. For each inhibitor, we compute the number of contacts between each heavy atom of the ligand and the center of mass of the side chain of each amino acid. A contact is defined to occur when that distance is less than 6 Å (see Section SIII of the SM for methodological details). Such threshold is chosen to capture specific non-covalent interactions while ensuring meaningful intermolecular binding^76^ (see Figure S3 of the SM for further detailed protein–ligand contact frequency maps). The number of contacts per heavy atom is normalized by the highest frequency for each ligand to allow comparison of the relative distribution of contacts across all inhibitors studied for both PARP1 (Figures 3A–D) and PARP2 (Figures 3E–H). Consequently, the relative contact score, ranging from 0 to 100 (%), is represented on a rainbow color scale. A summary of the most relevant protein residues which contact each ligand is provided in Table IA for PARP1 and Table IB for PARP2. In particular, only interactions that persisted over 80% of the total simulation time are included, ensuring that the reported contacts are both stable and functionally relevant.

**FIG. 3.**
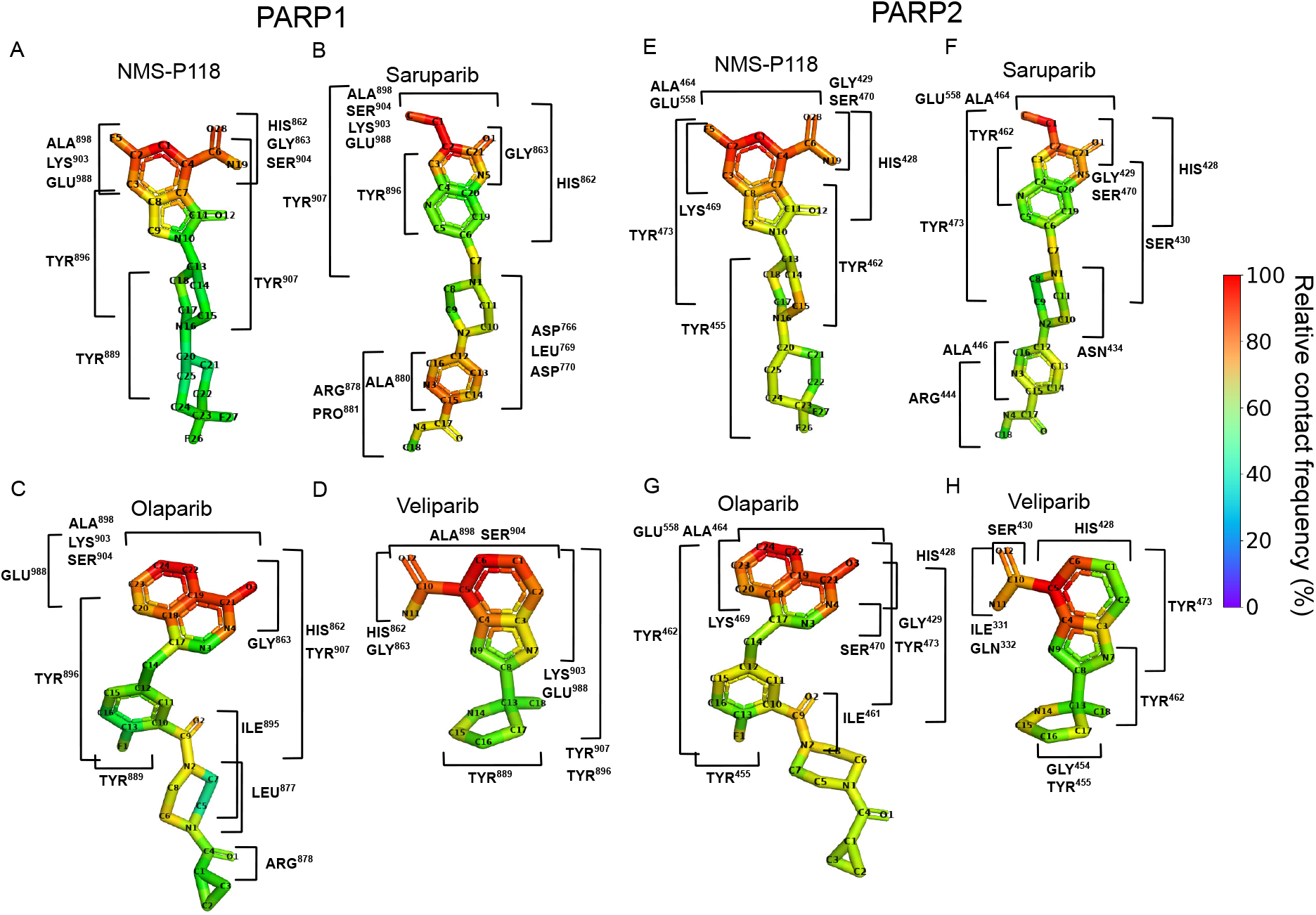
Relative contact frequency (%) of the PARP inhibitors with PARP1 (saruparib (A), NMS-P118 (B), olaparib (C), and veliparib (D)) and with PARP2 (saruparib (E), NMS-P118 (F), olaparib (G), and veliparib (H)) within the corresponding catalytic binding site. The relative contact frequency was calculated as the number of contacts per heavy atom, normalized by the highest value for each ligand-protein complex.

In Figure 3A, we present the normalized contact frequency of NMS-P118 within the catalytic binding pocket of PARP1. The atoms exhibiting the highest contact frequency are located within the fluorinated aromatic group (C2–C1–C4) and the amide moiety (C6–O28–N19). The fluorinated ring predominantly interacts with LYS903, consistent with the positive charge of the lysine side chain. The amide moiety also engages in multiple polar interactions with SER904, HIS862, and GLY863.

Moreover, NMS-P118 forms *π*–*π* stacking interactions with several tyrosine residues (TYR889, TYR896, and TYR907), which likely contribute to the ligand’s residence time within the active site and to the cooperative stabilization of the contact network governing its binding affinity. Notably, residues HIS862, GLY863, TYR889, TYR896, ALA898, LYS903, SER904, and TYR907 maintained contacts for more than 80% of the total simulation time (see Table IA), in agreement with static X-ray crystallographic structures of the corresponding lig- and–protein complexes^38^. This consistency supports the contact network predictions derived from our MD simulations.

The contact frequency analysis between PARP1 and saruparib is shown in Figure 3B. In this case, the region of the ligand exhibiting the highest frequency of contacts is centered around the aromatic ring at the lower part of the molecule (C12–C16 and N3). The most frequent and persistent interactions within the catalytic pocket involve the same residues observed for NMS-P118—namely HIS862, GLY863, ALA898, LYS903, SER904, TYR896, and TYR907—suggesting a conserved binding mode driven by a shared network of key non-bonded interactions despite the structural differences among both ligands (see Table IA). The tail region of saruparib displays non-bonded interaction modes between the amino groups of the non-aromatic ring and the residue side chains of ASP766 and ASP770. Moreover, the terminal amide group (centered on C17) engages in strong electrostatic interactions with ARG878, further stabilizing the ligand within the binding pocket. Importantly, the interactions identified from our simulations are consistent with those observed in X-ray crystallographic structures— ASP770, HIS862, GLY863, ARG878, ILE879, SER904, and TYR907—of the PARP1–saruparib complex^37^. Furthermore, our simulations reveal some transient additional contacts arising from the intrinsic flexibility of both the protein and the ligand, such as ASP776, LEU769, ALA880, PRO881, TYR896, ALA898, LYS903 and GLU988, which are not reported in static crystallographic configurations.

In Figure 3C, we present the interaction profile between olaparib and the catalytic domain of PARP1. The overall contact pattern closely resembles that of saruparib, reflecting the structural similarities between the two molecules, including their shared interaction with ARG878 (see Table IA and Figures 3A,C), but with considerably lower frequency in olaparib, hinting a potential mode of selectivity. Moreover, olaparib displays specific interactions including *π*–*π* stacking interaction with TYR889 through its fluorinated aromatic ring (C12–C13–F1), similar to that observed for NMS-P118 but with lower persistence (see the second row in Table IA). Interestingly, LEU877 and ILE895 emerge as unique contacts for olaparib, promoting its binding through non-polar interactions specific to its chemical structure. Although X-ray crystallography has resolved several of the key interactions between PARP1 and olaparib—particularly those involving catalytic residues^27^ (GLY863 and TYR907)—our atomistic MD simulations provide an extended comprehensive depiction of the binding interface including additional interactions such as HIS862, TYR889, ALA898, TYR896, LYS903, SER904, and GLU988.

Finally, the contact profile between veliparib and PARP1 is given in Figure 3D. The binding mode of veliparib is characterized by extensive *π*–*π* stacking interactions involving the aromatic residues TYR889, TYR896, and TYR907, distributed along a significant region of the ligand due to its smaller molecular size compared to the other inhibitors. Additionally, our results reveal that the most frequent contacts are concentrated around the core of the ligand (i.e., the bicyclic aromatic rings, C1–C6, and the C10-centered amide), corresponding to its most reactive region. The persistent nature of these core interactions, together with the distributed *π*–*π* stacking, likely enhances the overall affinity, and anchors veliparib within the active site.

Similarly, in Figure 3E–H we present the contact analysis between PARP2 and the different inhibitors: NMS-P118 (E), saruparib (F), olaparib (G), and veliparib (H). Overall, the interaction profiles resemble those observed for PARP1, involving comparable types of non-covalent contacts with homologous aromatic and polar residues within the binding pocket. Notably, when focusing on selective ligands (i.e., NMS-P118 and saruparib), their interaction distributions appear more homogeneous in PARP2. This effect likely reflects intrinsic differences in the binding interface, which in PARP2 are slightly more extended and permissive compared to PARP1, thereby enabling milder interactions with the specific regions of the ligands. For all PARP2–ligand complexes except for NMS-P118, the maximum contact frequency is lower than in their corresponding PARP1 complexes. However, in the case of NMS-P118, a broader contact distribution is maintained, suggesting genuine structural differences in ligand engagement that may contribute to the binding selectivity. In saruparib, the contacts involving the pyridinecarboxamide moiety—less buried within the active site—are markedly reduced in PARP2 compared to PARP1. Thus, while these four inhibitors display broadly similar binding modes between PARP1 and PARP2 at the residue identity level, the observed variations in contact distribution, absolute frequency, and spatial interconnectivity indicate a crucial role of the contacts for ligand recognition, with selectivity arising from subtle energetic differences in how these interaction networks are organized between both proteins.

**TABLE 1.**
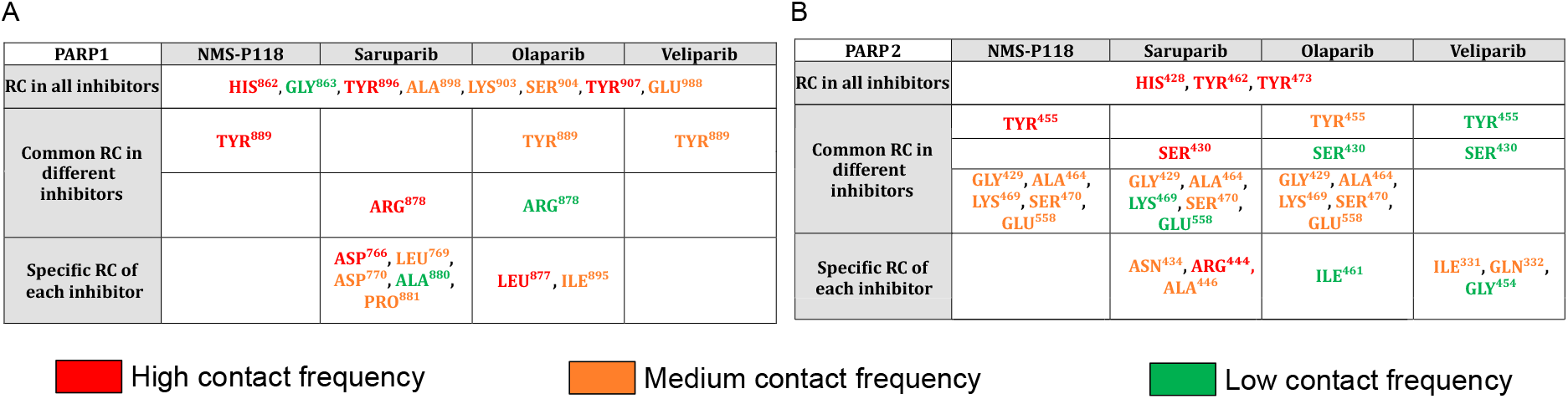
Key protein residues contacting (RC) the different inhibitors forming complexes with PARP1 (A) and PARP2 (B). Residues are colored according to the number of heavy-atom contacts established with the ligand: red indicates residues forming more than six contacts, orange corresponds to residues forming three to six contacts, and green highlights those forming one to three contacts. Rows indicate residue categories: contact residues present in all complexes, common residues among different inhibitors, and specific residue contacts of each inhibitor.

Analogously, Table IB lists the residues that consistently interact with all inhibitors in complex with PARP2. In contrast to PARP1, only a limited set of residues—specifically HIS428, TYR462, and TYR473— form common contacts across all ligands, suggesting a more restricted conserved interaction core within the PARP2 binding pocket at the level of residue identity. As with the PARP1 analysis, grouping the ligands according to their PARP1 selectivity does not reveal nitid contact patterns at the level of individual residue identity that are exclusive to either selective or non-selective compounds. Nonetheless, certain trends emerge within different subgroups. For instance, NMS-P118 and the non-selective inhibitors all interact with TYR455, while saruparib and the non-selective compounds share contacts with SER430, although with different binding probabilities. Interestingly, a broader set of shared contacts is observed between the PARP1-selective inhibitors and olaparib, including residues such as GLY429, ALA464, LYS469, SER470, and GLU558. This overlap suggests that olaparib may partially mimic the interaction profile and contact connectivity of PARP1-selective inhibitors within PARP2, potentially explaining its relatively balanced affinity for both protein paralogs. In general, our contact analysis captures ligand-specific patterns in residue interactions and binding poses, which together with the PMF free energy analyses provide a comprehensive view of the molecular determinants of PARP1 selectivity.

### Binding Interaction Analysis of PARP Inhibitors

To further characterize the nature of the protein–ligand interactions, we next carry out a per-residue binding energy decomposition for PARP1, aimed at identifying the dominant stabilizing forces and comparing how different inhibitors engage the catalytic pocket. This analysis resolves the specific energetic contributions from van der Waals (vdW), electrostatic (El), and solvation terms, the latter separated into polar solvent (PS) and non-polar solvent (NPS) components, providing a quantitative description of the energetic basis underlying inhibitor binding. The decomposition is carried out using the Molecular Mechanics/Poisson-Boltzmann Surface Area (MM/PBSA) approach (see Section IV of the SM for further technical details). Our results are summarized in Figure 4, which depicts both the spatial distribution of key energetic contributors mapped onto the protein surface (left panels) and the residue-specific energy components (right panels).

**FIG. 4.**
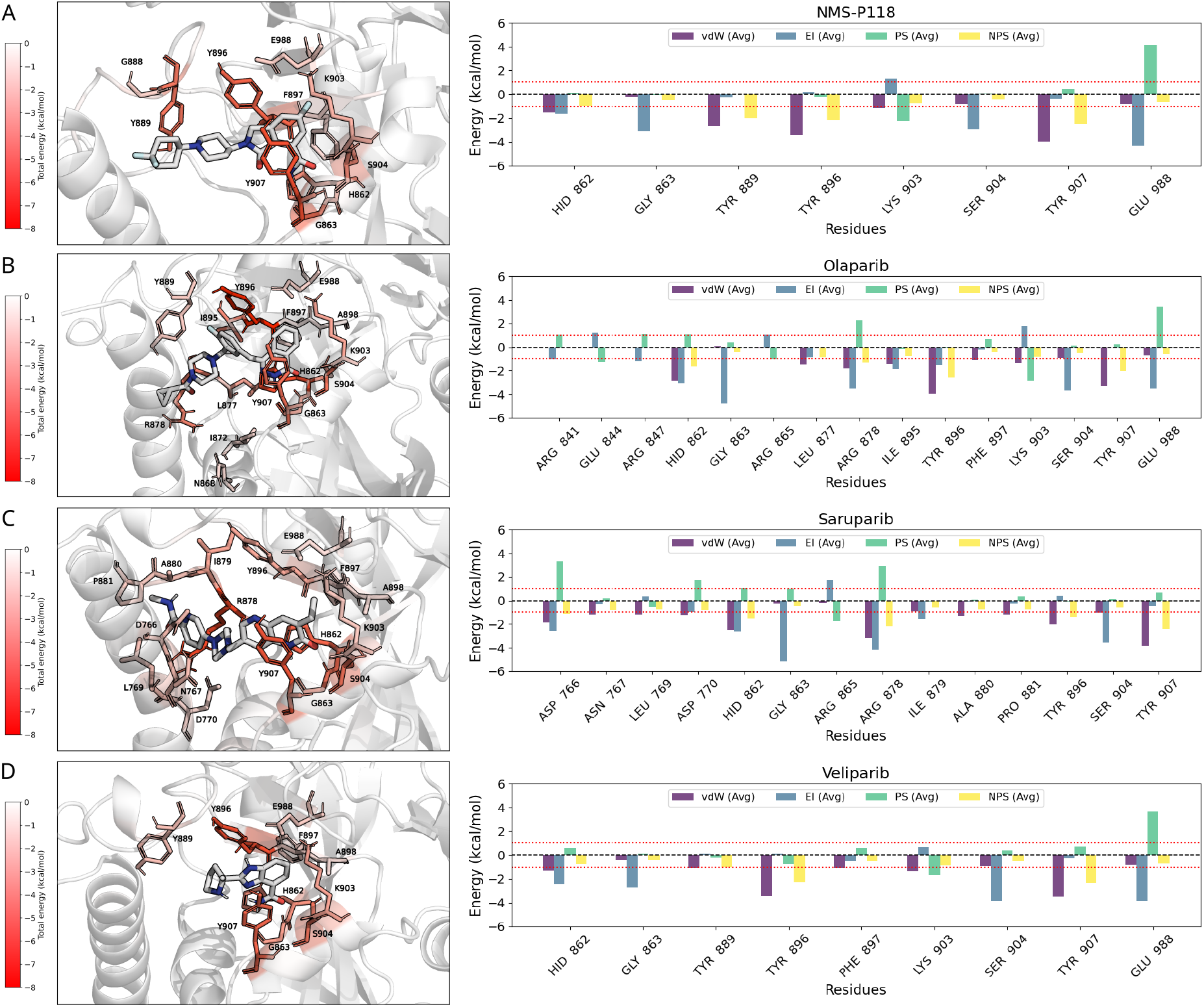
Binding-site energy decomposition for PARP1 in complex with NMS-P118 (A), olaparib (B), saruparib (C), and veliparib (D). The inhibitors are depicted in white while the identified interactions of the residues through a color map. Left panels: per-residue binding free energy decomposition mapped onto the protein surface, colored from low (white) to high (red) energetic contribution (kcal/mol). Right panels: average residue-wise interaction energies between PARP1 and each inhibitor. Only residues with an absolute contribution greater than 1 kcal/mol are displayed (depicted by the dashed red line). Energy components are decomposed into van der Waals (vdW, blue), electrostatic (El, purple), polar solvation (PS, green), and non-polar solvation (NPS, yellow) terms. Negative energy values referred to stabilizing ligand-protein residue contacts while positive values to destabilizing cross-interactions.

For the NMS-P118-PARP1 complex (Figure 4A), the strongest contributions arise from the aromatic residues TYR889, TYR896, and TYR907, primarily through van der Waals interactions, complemented by electrostatic stabilization from HIS862, GLY863, and GLU988. LYS903 also contributes through polar solvation effects that help secure the ligand within the catalytic site. In the olaparib-PARP1 complex (Figure 4B), the same tyrosine triad (TYR889, TYR896, TYR907) again dominates, although additional stabilization arises from ARG847 and GLU988, reflecting the distinct geometry of the olaparib scaffold. Saruparib (Figure 4C) engages the same canonical tyrosine residues, but a broader network of interactions is observed, with ASP766, ASN767, and LEU769 also providing stabilizing contributions, consistent with its extended contact surface within the binding pocket. Finally, veliparib (Figure 4D) maintains the general pattern of tyrosine-centered van der Waals stabilization, while electrostatic contributions from HIS862 and polar solvation effects from LYS903 are comparatively more pronounced, balancing its overall energetic signature. Across all complexes, van der Waals interactions consistently constitute the dominant stabilizing contribution, supported by residue-specific electrostatic contributions and, to a lesser extent, by polar solvation. In contrast, non-polar solvation contributions are marginal in multiple cases, with values generally close to the threshold of 1 kcal/mol. This relatively uniform energetic profile suggests that, within the resolution of the MM/PBSA framework, PARP1–ligand binding is predominantly stabilized by van der Waals and electrostatic interactions, as well as by localized polar contacts, whereas non-polar solvation contributions appear comparatively negligible. Nevertheless, subtle differences in residue-level contributions—such as the broader stabilization network in saruparib or the enhanced polar solvation terms in veliparib—highlight how small variations in chemical structure can fine-tune the energetic landscape of ligand binding.

The same binding free energy decomposition is performed for PARP2 (see Figure S4). The average binding energies for all inhibitors are shown in Figures S5A and S6A for PARP1 and PARP2 respectively, while the residue-level energetic contributions are detailed in Figures S5B-D and S6B-D. Our analysis indicates that the overall energetic landscapes of PARP1 and PARP2 are alike at the level of residue-resolved MM/PBSA decomposition. In both proteins, the dominant stabilizing forces arise from van der Waals interactions, primarily mediated by the conserved tyrosine triad (TYR889, TYR896, and TYR907 in PARP1; TYR462, TYR473,

and TYR455 in PARP2), together with secondary contributions from HIS862, GLY863, and GLU988 (HIS428, GLY429, and GLU558 in PARP2). Polar solvation terms associated with LYS903 (LYS469 in PARP2) also provide measurable stabilization, whereas non-polar solvation contributions remain marginal in all cases. When comparing distinct inhibitors, no pronounced differences are detected in the average interaction energies or in the residue-specific decomposition profiles within the statistical resolution of the MM/PBSA analysis (Figures S5 and S6), or in the hydrogen bond analysis (Figure S7). Both selective and non-selective inhibitors engage a largely overlapping set of residues with broadly comparable energetic contributions, and the relative weighting between van der Waals and electrostatic components remains similar across ligands at this level of analysis. This outcome suggests that residue-level binding free energy analysis through MM/PBSA, while highly informative for identifying residues that stabilize the complex, might be insufficient by itself to fully resolve the molecular origin of binding selectivity between closely related paralogs.

### Impact of Single-point Mutations in PARP1-saruparib Binding Interaction Energy

To gain deeper insight into the residue-level determinants of selective inhibitors of PARP1 such as saruparib, we now apply a systematic mutational analysis targeting key amino acids within the active site, previously identified as major interactions in the bound-state (Figs. 3 and 4). We have selected saruparib for these calculations over NMS-P118 not only due to its larger, experimentally reported selectivity^41^, but also because it exhibits several distinctive features: (1) it shows the most pronounced difference in PMF profiles between PARP1 and PARP2 (Fig. 1D); (2) it retains a high degree of PARP1 conformational flexibility (see Section V and Figures S8C, S8H of the SM for further details); and (3) it forms a broad set of intermolecular contacts (Fig. IA), particularly involving its core and tail regions. These features make saruparib the most informative model among the different tested inhibitors to dissect the residue specific contributions of ligand binding. The mutated residues are selected based on contact frequency—specifically those within the active site forming interactions with the largest number of heavy atoms in saruparib. This criterion provides a complementary view of binding determinants, as mutating these residues to glycine is expected to simultaneously disrupt multiple protein–ligand contacts, thereby revealing the extent to which binding stability depends on an intricate network of side-chain interactions. In particular, residues are chosen according to the number of contacts established with saruparib when measured from their C*_α_* atom within a 6 Å cut-off distance, offering an alternative perspective to that presented in Figure 3. By replacing the selected residues with glycine, side-chain contributions are minimized, allowing the assessment of their roles in maintaining the integrity of the ligand-protein binding. The targeted residues—LEU877, ARG878, ILE879, ILE895, and ASN906—were identified as major contributors, among few others, to saruparib PARP1-binding. To systematically evaluate their energetic relevance, different mutation sets are generated, comprising simultaneous replacements of five, three, or two amino acids, thereby enabling a graded assessment of their collective impact on ligand stabilization. Contact maps of the active-site region for the wild-type and mutant complexes (Figures 5A–D) reveal pronounced alterations in the protein–ligand interaction network. In the wild-type PARP1–saruparib complex (Figure 5A), an extensive network of close, highly frequent contacts is observed, reflecting multiple cooperative stabilizing interactions between the ligand and the surrounding protein residues within the pocket.

**FIG. 5.**
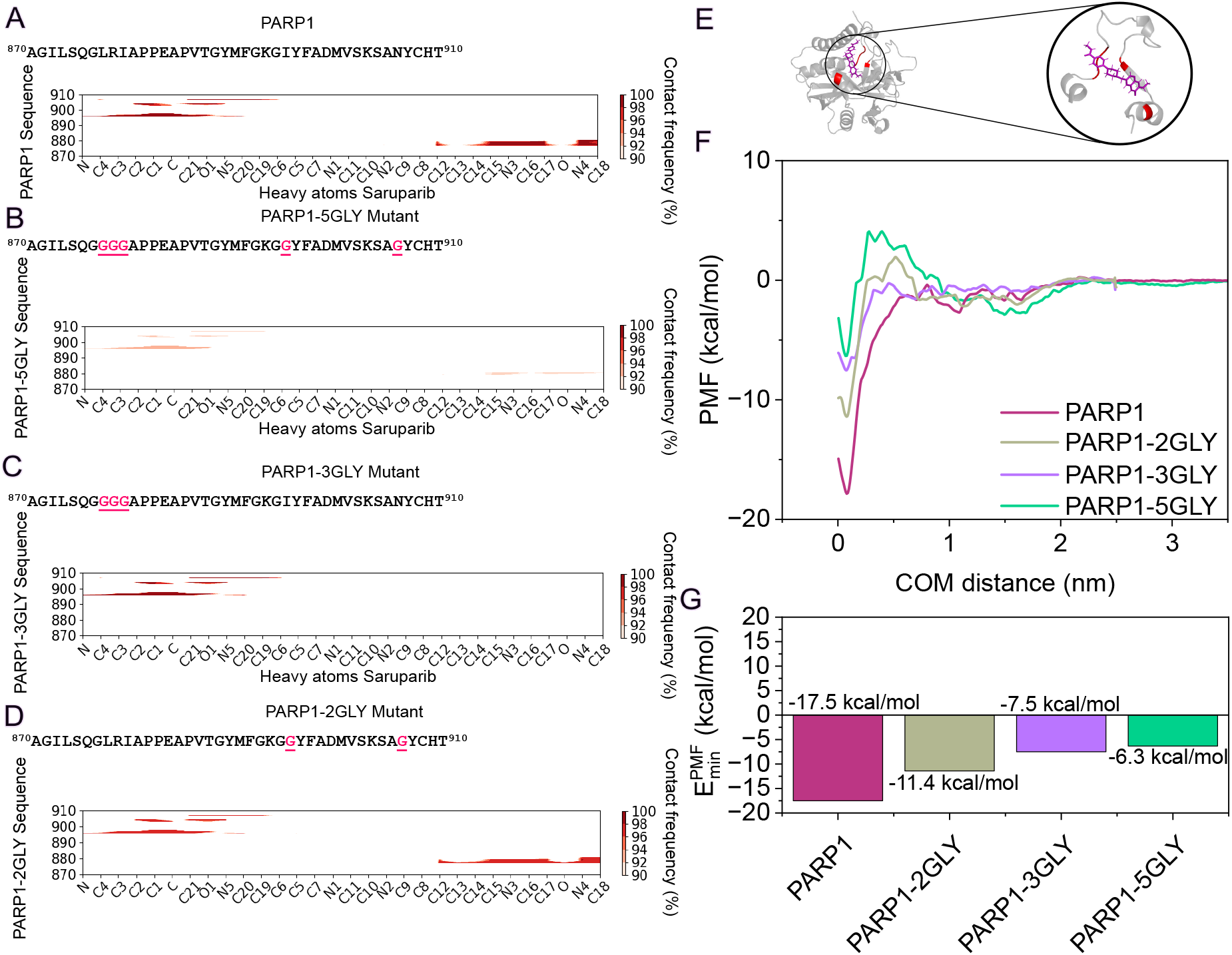
Residue–ligand contact frequency maps for saruparib bound to PARP1 (A) and its glycine-substituted variants: 5GLY (B), 3GLY (C), and 2GLY (D). The PARP1 sequence region corresponding to residues 870–910 is shown above each map, with glycine substitution sites highlighted in red. Contact frequencies are calculated between the heavy atoms of saruparib and the protein residues within the active site. (E) Cartoon representation of the PARP1 active site in complex with saruparib highlighting the mutated amino acids in red for the 5GLY mutant. (F) PMF profiles for saruparib-PARP1 complex and the glycine mutants (PARP1-2GLY, PARP1-3GLY, and PARP1-5GLY), computed as a function of the center-of-mass distance between the ligand and the protein binding pocket. (G) Comparison of the minimum binding free energy values obtained from the PMF curves in (F).

Mutation of five key residues to glycine (PARP1-5GLY mutant: LEU877, ARG878, ILE879, ILE895, and ASN906; Figure 5B) leads to a substantial reduction in ligand-protein contacts across the catalytic site, particularly affecting both the 877–879 segment and the 895–906 region. In the triple glycine mutant (PARP1-3GLY: LEU877, ARG878, ILE879; Figure 5C), the loss of contacts within the 877–879 region is retained, while interactions between residues 895–906 are largely preserved. This indicates that the main hotspot for stabilizing interactions is within the 877–879 segment. The double glycine mutant (PARP1-2GLY: ILE895 and ASN906; Figure 5D) exhibits only a moderate reduction in interaction intensity, while the overall contact pattern remains largely comparable to that of the wild-type complex (Figure 5A). The structural visualization of the catalytic site (Figure 5E) highlights the mutated residues in red, illustrating their spatial arrangement within the binding cleft and their contribution to shaping the complementary surface accommodating saruparib. Collectively, these analyses suggest that while the binding interface is broadly distributed across the catalytic pocket, the 877–879 segment represents a major hotspot for stabilizing ligand interactions, whereas other residues primarily modulate interaction strength without markedly altering the overall intermolecular contact landscape and binding pose.

Next, we quantify the variations in the molecular interaction network found across different contact maps through PMF calculations, providing a detailed energetic description of the binding interaction strength for the wild-type *vs.* mutated PARP1 sequences in complex with saruparib. The resulting PMF profiles, depicted in Figure 5F, and their corresponding minimum values in Figure 5G, reveal a systematic modulation of the binding free energy landscape as a function of the number of glycine substitutions. The wild-type PARP1 complex (maroon curve) is characterized by a deep, well-defined global minimum of –17.5 kcal/mol, reflecting a highly stable interaction driven by optimal complementarity between the ligand and the intact active site. The profile displays a smooth descent into the energy well, indicative of a smooth pathway for ligand association.

We find that while the introduction of mutations progressively diminishes the binding stability, the effect is not entirely uniform across the different mutants. The double glycine mutant (PARP1-2GLY; grey curve) exhibits a well-defined minimum of –11.4 kcal/mol— shallower than the wild-type but still clearly favorable. Nevertheless, the reduction in depth by approximately 6 kcal/mol highlights the importance of these two residues controlling the binding affinity. Notably, the emergence of more pronounced energy barriers at intermediate distances suggests that the mutated residues may also play a role in favoring the ligand pathway into the active site. In contrast, the triple mutant (PARP1-3GLY, violet curve) displays a less stable minimum of –7.5 kcal/mol, reflecting the energetic contributions of the additional side chains to the final bound state. In this case, the dominant effect is not the emergence of intermediate barriers but rather a destabilization of the binding minimum itself, underscoring the significant role of these three residues in maintaining the thermodynamic stability of the complex. Finally, the most severe energetic perturbation is observed for the quintuple mutant (PARP1-5GLY, green curve), where both effects combine: the minimum rises further to –6.3 kcal/mol, and the PMF profile shows a more pronounced energy barrier of 4.5 kcal/mol preceding it. This indicates that the extensive loss of complementary interactions simultaneously weakens the final bound state and introduces significant hurdles to ligand attachment, likely associated with desolvation penalties triggered by the absence of stabilizing protein contacts.

Collectively, our PMF calculations indicate that the high-affinity binding of saruparib to PARP1 cannot be attributed to a single dominant interaction, but instead emerges from a cooperative network of multiple residue contacts. This conclusion is supported by the observation that mutating a single residue (ARG878) produces only localized changes in the interaction network and results in a modest perturbation of the overall protein–ligand binding free energy (Figure S9 of the SM), although it may be crucial for binding selectivity^55^ (see Table I). In contrast, simultaneous mutation of several key residues to glycine leads to cumulative perturbations of the interaction network, resulting in a measurable impact on the thermodynamics of binding. This detailed energetic mapping provides a molecular framework for interpreting structure–activity relationships and offers mechanistic guidance for the rational design of inhibitors with enhanced selectivity. For completeness, the full set of contact maps for the mutant complexes is presented in Figure S10 of the SM.

## CONCLUSIONS

In this work, we present an atomistic study of how four representative PARP inhibitors engage PARP1 and PARP2, and how these interactions collectively contribute to binding selectivity. By performing atomistic MD simulations and computing contact frequency maps, energy decomposition analyses, umbrella sampling potential-of-mean-force calculations, and targeted mutational perturbations, we demonstrate that selectivity is driven by a cooperative multivalent network of stabilizing contacts anchoring the ligand within the pocket. Such network of intermolecular contacts governs the thermodynamic stability and binding free energy of the ligand-protein complex.

PMF calculations help clarify the mechanistic basis of selectivity (Figure 1). We find that selective inhibitors such as NMS-P118 and Saruparib show the largest differences in binding free energy between PARP1 and PARP2, displaying binding landscapes that not only favor the formation of the desired complex but also present energetic barriers hindering association with the non-target paralog, whereas non-selective compounds exhibit smoother energy surfaces that facilitate association with both proteins. In other words, specificity can be partially regulated by the association energy pathway rather than solely in the thermodynamic stability of the end-point. Notably, comparison of PMF-derived Δ*G* values with experimental estimates obtained from IC_50_ measurements reveals a remarkable correlation, indicating that simulations reproduce the relative selectivity of the inhibitors despite systematic offsets in absolute free energies (Figure 2).

Our contact frequency analysis provide microscopic insights into the binding mechanism in agreement with experimental observations, highlighting the central amino acid–ligand interactions most critical for stable complex formation. The most frequent contacts are notably preserved between PARP1 and PARP2, and are primarily mediated by a tyrosine triad through van der Waals interactions (TYR889, TYR896, TYR907 in PARP1; TYR455, TYR462, TYR473 in PARP2). Contact frequency and energetic decomposition analyses reveal a conserved network of stabilizing interactions that defines the bound-state pose within the catalytic pocket. These common interactions are necessary for binding to the active site, while selectivity arises from how these contacts are energetically weighted, and how other contributing interactions—mostly electrostatic— cooperatively emerge to define the protein-ligand binding free energy (Figures 3, 4).

The mechanistic picture given by the molecular contact analysis and the thermodynamic evaluation provided by PMF calculations with increasing number of residue mutations have direct implications in binding affinity. Maximizing selectivity requires more than lowering the overall binding free energy, instead relying on design choices that enhance interactions with structural features surrounding the binding pocket, stabilize the desired bound conformation, and suppress off-target binding. In that sense, identifying the key contacts driving ligand-complex stabilization—and how these contacts cooperatively reshape the binding association pathway—is fundamental to maximizing specific binding. Our mutational perturbation analysis demonstrates that selective binding relies on multivalent cooperative interactions, rather than on single dominant interaction modes (Figure 5). Specifically, simultaneous mutation of five key amino acids to glycine within the binding interface reduces binding affinity by more than 50%, thereby preventing stable complex formation. Therefore, designing ligands that engage multiple complementary motifs— e.g., multivalent—likely yields more durable protein-specific inhibitors than ligands relying on a highly specific binding mode.

Taken together, our study highlights PARP inhibitor selectivity as an energy landscape problem: the critical question is not simply how tightly a ligand binds, but how the entire route to binding is shaped and modulated by a cooperative network of ligand–protein interactions. By computing contact molecular networks and potential-of-mean-force calculations, we propose a computational framework to guide and rationalize the design of next-generation enzyme-selective inhibitors with improved efficacy and safety.

## Data availability

We provide the relevant data in the repository (GitHub link for the repository) to facilitate reproducibility of our results. In the repository we also give the necessary code to run the simulations and accessible instructions to obtain our results.

## ACKNOWLEDGMENTS

A. F. acknowledges funding from the Ramon y Cajal fellowship (RYC2021-030937-I) and Spanish National Grant (PID2022-136919NA-C33). A. R. T. acknowledges funding from the Juan de la Cierva fellowship (JDC2024-053759-I). J. R. E. acknowledges funding from Emmanuel College, the University of Cambridge, the Ramon y Cajal fellowship (RYC2021-030937-I), the Spanish scientific plan and committee for research reference PID2022-136919NA-C33, and the European Research Council (ERC) under the European Union’s Horizon Europe research and innovation program (grant agreement no. 101160499). A.O., N.D.-V, C.P. and L.P.-H. acknowledge funding from CRIS Cancer Foundation (AOF.C01CRIS and AOF.M01CRIS). This work has also been performed using resources provided by Archer2 (https://www.archer2.ac.uk/) funded by EPSRC Tier-2 capital grant EP/P020259/e829. The authors also thankfully acknowledge RES computational resources provided by Mare Nostrum 5 through the activities 2024-3-0001 and 2025-1-0009.

