## Supplementary Material for "Cooperative molecular interaction networks govern PARP1 inhibitor selectivity and binding affinity"

Alejandro Feito

*Department of Physical Chemistry, Universidad Complutense de Madrid, Av. Complutense s/n, Madrid 28040, Spain and Instituto Pluridisciplinar, Universidad Complutense de Madrid, P.º de Juan XXIII, 1, Moncloa - Aravaca, 28040 Madrid, Spain*

Natàlia DeMoya-Valenzuela, Cristian Privat, and Lucía Paniagua-Herranz

*Experimental Therapeutics Unit, Hospital Clínico San Carlos (HCSC) Instituto de Investigación Sanitaria San Carlos (IdISSC), Madrid, Spain*

Andrés R. Tejedor

*Department of Physical Chemistry, Universidad Complutense de Madrid, Av. Complutense s/n, Madrid 28040, Spain Yusuf Hamied Department of Chemistry, University of Cambridge, Lensfield Road, Cambridge CB2 1EW, UK and Instituto Pluridisciplinar, Universidad Complutense de Madrid, P.º de Juan XXIII, 1, Moncloa - Aravaca, 28040 Madrid, Spain*

Adiran Garaizar

*Data Science, Bayer AG, Alfred-Nobel-Straße 50, 40789 Monheim am Rhein, Germany*

Alberto Ocana

*Experimental Therapeutics Unit, Hospital Clínico San Carlos (HCSC) Instituto de Investigación Sanitaria San Carlos (IdISSC), Madrid, Spain and PhAsIca Biosciences S.L, Calle Velázquez, 27, 28001 Madrid, Spain*

Jorge R. Espinosa

*Department of Physical Chemistry, Universidad Complutense de Madrid, Av. Complutense s/n, Madrid 28040, Spain*

*Yusuf Hamied Department of Chemistry, University of Cambridge,*
*Lensfield Road, Cambridge CB2 1EW, UK*
*Instituto Pluridisciplinar, Universidad Complutense de Madrid,*
*P.º de Juan XXIII, 1, Moncloa - Aravaca, 28040 Madrid, Spain and*
*PhAsIca Biosciences S.L, Calle Velázquez, 27, 28001 Madrid, Spain*
(Dated: January 16, 2026)

### SI. PREPARATION OF THE SYSTEMS AND SIMULATION DETAILS

All atomistic simulations were performed using the GROMACS simulation package (version 2023)[1]. Given our interest in exploring interaction potentials at physiological salt concentration (150 mM NaCl), with a solvent and ion model that accurately reproduces ion solubilities in water at 310 K, these configurations were then solvated in a cubic box ($8.5 \times 8.5 \times 8.5$ nm; for those in Figure **??**, the box size was $10 \times 10 \times 10$ nm). Some water molecules were replaced by $Na^+$ and $Cl^-$ ions to neutralize the system and achieve the desired salt concentration. Energy minimization was performed with a force tolerance of 1000 kJ $mol^{-1}$ $nm^{-1}$. Bond constraints on hydrogen-containing bonds were enforced via the LINCS algorithm [2], allowing a time step of 2 fs. Periodic boundary conditions (PBC) were maintained, and electrostatics were computed using the Particle-Mesh Ewald (PME) [3] method with a Coulomb cutoff of 0.9 nm. Temperature coupling was performed using the velocity-rescale (v-rescale) thermostat with a relaxation time constant $\tau_T = 1.0$ ps, while pressure coupling was applied using the Parrinello–Rahman barostat with a relaxation time constant $\tau_P = 1.0$ ps.

For the PMF simulations the configurations were solvated in a box of 8.5x8.5x15 nm at 300K and 150mM of NaCl. The center-of-mass (COM) distance between the protein and the ligand controlled using a harmonic umbrella potential with a pulling force constant of 10000 kJ $mol^{-1}$ $nm^{-2}$. Approximately 60 umbrella sampling windows were defined, spaced every 0.025 nm within the range. For production runs, positional restraints of 1000 kJ $mol^{-1}$ $nm^{-2}$ (in directions perpendicular to the pulling axis) were applied to the amino acid heavy atoms of four residues of the protein with the most values of frequency contacts along the simulation with the ligand (to avoid rotations) and the heavy atoms of the drug were placed at the center of the molecule to allow freedom of movement of the regions that predominantly interact with the protein during the simulation. Each system was simulated for a total accumulated simulation time of 600 ns. The Weighted Histogram Analysis Method (WHAM) [4], as implemented in GROMACS, was used to reconstruct the free energy profiles. The initial 2000 ps of each simulation were excluded from WHAM analysis to ensure equilibration. Statistical errors of the PMF profiles were estimated by block analysis of the simulation trajectories. Each trajectory was divided into five independent segments, and PMF profiles were computed separately for each segment using the GROMACS WHAM tool. The statistical uncertainty at each point along the reaction coordinate was then quantified as the standard deviation of the PMF values obtained from these five blocks.

### SII. SEQUENCE OF PARP1 AND PARP2 AND PDB OF THE STRUCTURED DOMAINS AND SEQUENCE OF PARP1

PARP1:

MAESSDKLYRVEYAKSGRASCKKCSESIPKDSLRMAIMVQSPMFDGKVPHWYHFSCFWKVGHSIRHPDVEVDGFS
ELRWDDQQKVKKTAEAGGVTGKGQDGIGSKAEKTLGDFAAEYAKSNRSTCKGCMEKIEKGQVRLSKKMVDPEKPQ
LGMIDRWYHPGCFVKNREELGFRPEYSASQLKGFSLLATEDKEALKKQLPGVKSEGKRKGDEVDGVDEVAKKKSK
KEKDKDSKLEKALKAQNDLIWNIKDELKKVCSTNDLKELLIFNKQQVPSGESAILDRVADGMVFGALLPCEECSG
QLVFKSDAYYCTGDVTAWTKCMVKTQTPNRKEWVTPKEFREISYLKKLKVKKQDRIFPPETSASVAATPPPSTAS
APAAVNSSASADKPLSNMKILTLGKLSRNKDEVKAMIEKLGGKLTGTANKASLCISTKKEVEKMNKKMEEVKEAN
IRVVSEDFLQDVSASTKSLQELFLAHILSPWGAEVKAEPVEVVAPRGKSGAALSKKSKGQVKEEGINKSEKRMKL
TLKGGAAVDPDSGLEHSAHVLEKGGKVFSATLGLVDIVKGTNSYYKLQLLEDDKENRYWIFRSWGRVGTVIGSNK
LEQMPSKEDAIEHFMKLYEEKTGNAWHSKNFTKYPKKFYPLEIDYGQDEEAVKKLTVNPGTKSKLPKPVQDLIKM
IFDVESMKKAMVEYEIDLQKMPLGKLSKRQIQAAYSILSEVQQAVSQGSSDSQILDLSNRFYTLIPHDFGMKKPP
LLNNADSVQAKVEMLDNLLDIEVAYSLLRGGSDDSSKDPIDVNYEKLKTDIKVVDRDSEEAEIIRKYVKNTHATT
HNAYDLEVIDIFKIEREGECQRYKPFKQLHNRRLLWHGSRTTNFAGILSQGLRIAPPEAPVTGYMFGKGIYFADM
VSKSANYCHTSQGDPIGLILLGEVALGNMYELKHASHISKLPKGKHSVKGLGKTTPDPSANISLDGVDVPLGTGI
SSGVNDTSLLYNEYIVYDIAQVNLKYLLKLKFNFKTSLW

PARP1 Active Site:

NNADSVQAKVEMLDNLLDIEVAYSLLRGGSDDSSKDPIDVNYEKLKTDIKVVDRDSEEAEIIRKYVKNTHATTHN
AYDLEVIDIFKIEREGECQRYKPFKQLHNRRLLWHGSRTTNFAGILSQGLRIAPPEAPVTGYMFGKGIYFADMVS
KSANYCHTSQGDPIGLILLGEVALGNMYELKHASHISKLPKGKHSVKGLGKTTPDPSANISLDGVDVPLGTGISS
GVNDTSLLYNEYIVYDIAQVNLKYLLKLKFNFKTSLW

The following Protein Data Bank (PDB) codes were used for the atomistic simulations: PARP1 with saruparib (9ETQ [5]), with olaparib (7KK4 [6]), with veliparib (7KK6 [6]) and with NMS-P118 (5A00 [7]). For the wild-type protein we used the AlphaFold [8] prediction (model AF-P09874-F1).

PARP2:

MAARRRRSTGGGRARALNESKRVNNGNTAPEDSSPAKKTRRCQRQESKKMPVAGGKANKDRTEDKQDGMPGRSWA

SKRVSESVKALLLKGKAPVDPECTAKVGKAHVYCEGNDVYDVMLNQTNLQFNNNKYYLIQLLEDDAQRNFSVWMR
WGRVGKMGQHSLVACSGNLNKAKEIFQKKFLDKTKNNWEDREKFEKVPGKYDMLQMDYATNTQDEEETKKEESLK
SPLKPESQLDLRVQELIKLICNVQAMEEMMMEMKYNTKKAPLGKLTVAQIKAGYQSLKKIEDCIRAGQHGRALME
ACNEFYTRIPHDFGLRTPPLIRTQKELSEKIQLLEALGDIEIAIKLVKTELQSPEHPLDQHYRNLHCALRPLDHE
SYEFKVISQYLQSTHAPTHSDYTMTLLDLFEVEKDGEKEAFREDLHNRMLLWHGSRMSNWVGILSHGLRIAPPEA
PITGYMFGKGIYFADMSSKSANYCFASRLKNTGLLLLSEVALGQCNELLEANPKAEGLLQGKHSTKGLGKMAPSS
AHFVTLNGSTVPLGPASDTGILNPDGYTLNYNEYIVYNPNQVRMRYLLKVQFNFLQLW

PARP2 Active Site:

TQKELSEKIQLLEALGDIEIAIKLVKTELQSPEHPLDQHYRNLHCALRPLDHESYEFKVISQYLQSTHAPTHSDY
TMTLLDLFEVEKDGEKEAFREDLHNRMLLWHGSRMSNWVGILSHGLRIAPPEAPITGYMFGKGIYFADMSSKSAN
YCFASRLKNTGLLLLSEVALGQCNELLEANPKAEGLLQGKHSTKGLGKMAPSSAHFVTLNGSTVPLGPASDTGIL
NPDGYTLNYNEYIVYNPNQVRMRYLLKVQFNFLQLW

The following Protein Data Bank (PDB) codes were used for the atomistic simulations: PARP2 with olaparib (4TVJ [6]), with veliparib (3KJD [9]) and with NMS-P118 (4ZZY [7]). For the wild-type protein we used the AlphaFold prediction (model AF-Q9UGN5-F1).

### SIII. CALCULATING CONTACT MAPS

Computation of intermolecular and intramolecular contact maps within protein condensates is calculated from all-atom trajectories. Typically, molecular contacts are identified based on a distance criterion, with the assumption that the relative frequency of contact map occurrences (rather than absolute frequency) remains generally unaffected by the selected cut-off distance used in calculations, assuming the cut-off values are reasonable. In particular, we use a criterion that depends on the specific excluded volume of each amino acid, adopting a sequence-dependent cut-off distance equivalent to $1.2\sigma_{ij}$, where $\sigma_{ij}$ represents the mean excluded volume of the respective $i$th and $j$th amino acids [10]. For the heavy atoms of drug molecules, a $\sigma$ value of 5 Å was assigned. Given that the minimum of the potential used lies at approximately $2^{1/6}\sigma_{ij} \approx 1.122\sigma_{ij}$, we set the cut-off distance slightly beyond this point, at $1.2\sigma_{ij}$, to ensure significant binding.

### SIV. CALCULATING BINDING FREE ENERGY AND HYDROGEN BOND FORMATION

We estimated the binding free energy ($\Delta G_{binding}$) of the ligand-protein complexes using the MMPBSA.py module [11] from AmberTools23 [12] with the molecular mechanics generalized Born surface area (MMGBSA) approach. The binding free energy was calculated every 10 frames of the molecular dynamics simulation trajectory. The total $\Delta G_{binding}$ was decomposed into electrostatic, van der Waals (vdW), polar solvation, and non-polar solvation components (Eq. S11) through pairwise per-residue analysis (idecomp = 4).

$$\Delta G_{\text{binding}} = \Delta E_{\text{elec}} + \Delta E_{\text{vdW}} + \Delta G_{\text{polar}} + \Delta G_{\text{nonpolar}} - T\Delta S \tag{S1}$$

The electrostatic ($\Delta E_{elec}$) and van der Waals ($\Delta E_{vdW}$) terms were calculated directly from the molecular mechanics force field as the sum of pairwise atomic interactions. The polar solvation term ($\Delta G_{polar}$) was computed using the modified generalized Born model developed by Onufriev, Bashford, and Case (igb=5) [13], while the non-polar contribution ($\Delta G_{nonpolar}$) was estimated using solvent-accessible surface area (SASA) default model (gbsa=2). Default parameters were applied in both solvation contributions. The entropic component ($T\Delta S$) was not included in the estimation of the binding free energy.

In addition, hydrogen bond formation was analyzed throughout the trajectories using CPPTRAJ [14], enabling the identification of persistent hydrogen-bonding interactions between the ligand and protein residues. A hydrogen bond is defined as an interaction involving an acceptor heavy atom (A), a donor hydrogen atom (H), and its corresponding donor heavy atom (D). The interaction was identified present when the distance between the acceptor and donor heavy atoms was less than 3.0 Å and the A–H–D angle was greater than 135°.

### SV. CALCULATING ROOT MEAN SQUARE DEVIATION AND FLUCTUATION

We quantified the structural stability and flexibility of the PARP1 system by calculating the Root Mean Square Deviation (RMSD) and Root Mean Square Fluctuation (RMSF) from all-atom molecular dynamics trajectories using the package MDAnalysis in Python3.

The RMSD, which measures the average displacement of atoms relative to a reference structure over time, was calculated for the protein's C$\alpha$ atoms using the following formula:

$$\mathrm{RMSD}(t) = \sqrt{\frac{1}{N}\sum_{i=1}^{N} \left| r_i(t) - r_i^{\mathrm{ref}} \right|^2} \tag{S2}$$

where $N$ is the total number of atoms considered (in this case, C$\alpha$ atoms), $r_i(t)$ is the position vector of the $i$-th atom at time $t$, and $r_i^{\mathrm{ref}}$ is the position vector of the corresponding atom in the reference structure.

Subsequently, the RMSF was calculated to evaluate the per-residue flexibility. The RMSF for each residue $\alpha$ is defined as:

$$\mathrm{RMSF}(\alpha) = \sqrt{\frac{1}{T}\sum_{t=1}^{T} (r\alpha(t) - \langle r\alpha \rangle)^2} \tag{S3}$$

where $T$ is the total number of frames in the trajectory, $\vec{r}\alpha(t)$ is the position vector of the C$\alpha$ atom of residue $\alpha$ at time $t$, and $\langle \vec{r}\alpha \rangle$ is the time-averaged position of that same atom. For this analysis, the protein C$\alpha$ atom was first aligned to the first configuration of the simulation.

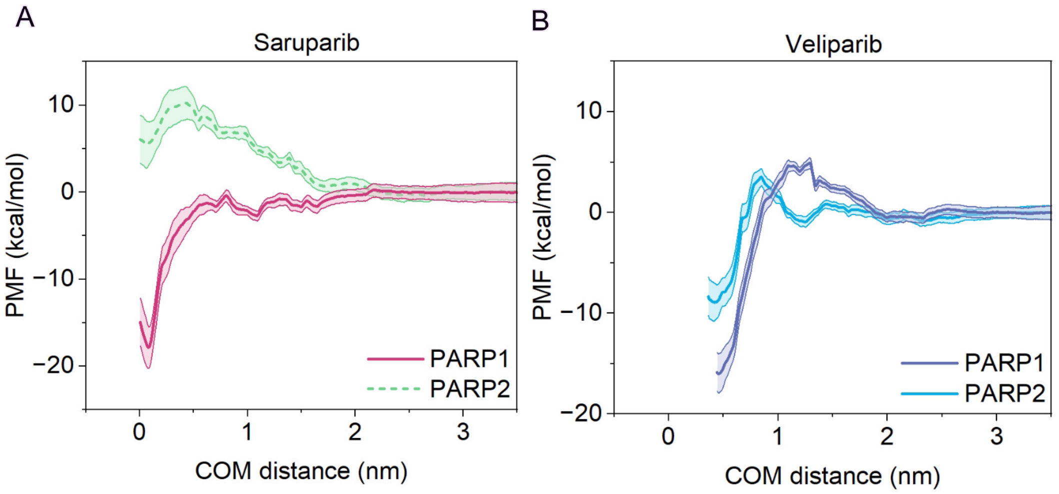


**FIG. S1:** Atomistic PMF dissociation profiles between each ligand and PARP1/PARP2 under physiological NaCl concentration (150 mM) and room conditions, in explicit solvent and ions. The COM distance between the protein binding pocket and the ligand is used as the reaction coordinate. Curves are shown for saruparib (A) and veliparib (B), depicting the statistical uncertainty of each PMF profile by a color shadowing obtained from bootstrapping analysis.

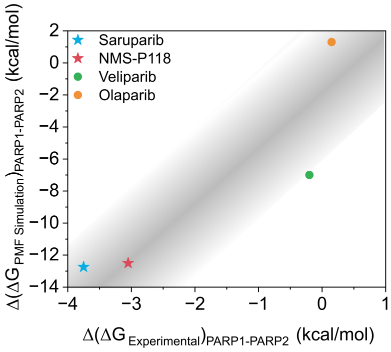


**FIG. S2:** Comparison between experimental ($\Delta(\Delta G_{\mathrm{Experimental}})_{\mathrm{PARP1-PARP2}}$) and simulated ($\Delta(\Delta G_{\mathrm{PMF\ Simulation}})_{\mathrm{PARP1-PARP2}}$) binding free energies for different inhibitors (Saruparib, NMS-P118, Veliparib, and Olaparib) for PARP1 and PARP2. A linear shading to the obtained correlation was added as a guide to the eye. Specific inhibitors of PARP1 are plotted in stars and non-specific inhibitors in circles.

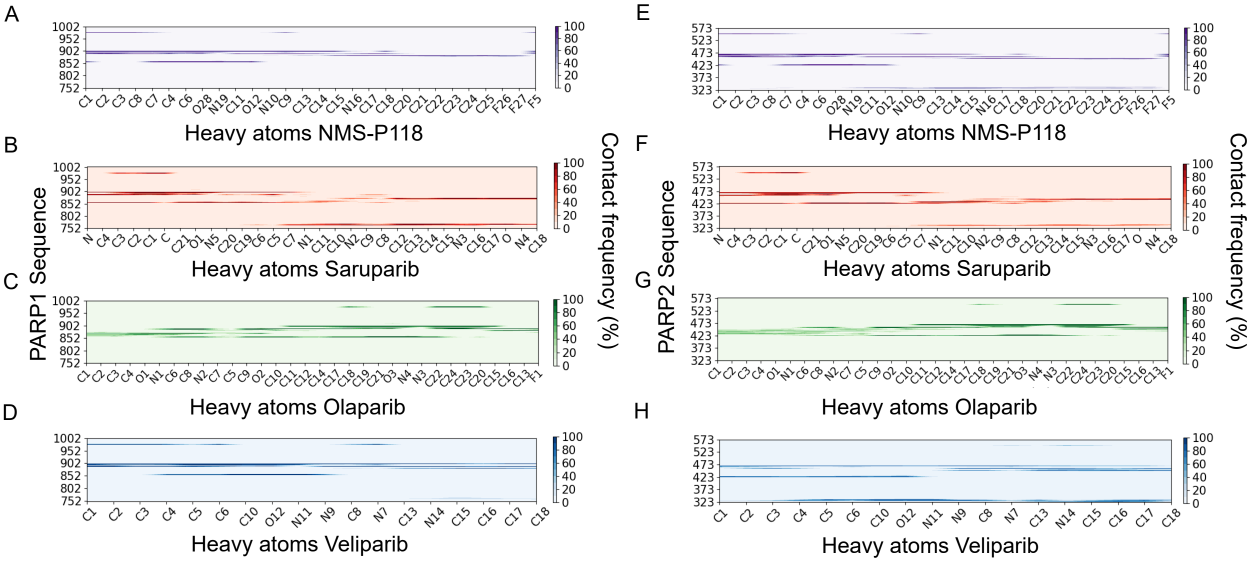


**FIG. S3:** Intermolecular frequency contact maps for PARP1 protein with heavy atoms of NMS-P118 (A), saruparib (B), olaparib (C) and veliparib (D). Intermolecular frequency contact maps for PARP2 protein with heavy atoms of NMS-P118 (E), saruparib (F), olaparib (G) and veliparib (H). The contacts are represented as contact frequency in percentage.

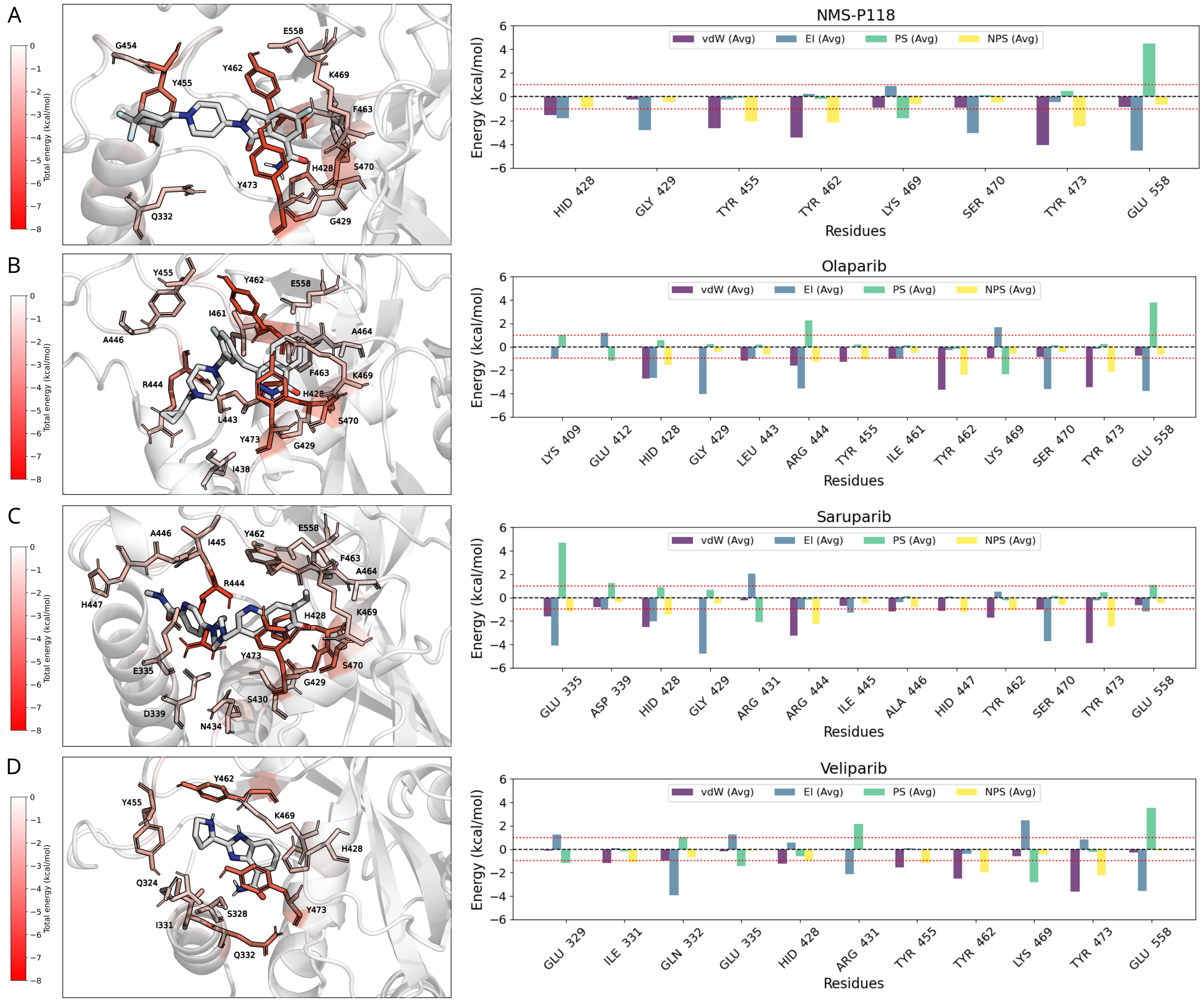


**FIG. S4:** Binding site interaction energies for PARP2 with NMS-P118 (A), olaparib (B), saruparib (C), and veliparib (D). Left panels: total per-residue binding free energy decomposition mapped onto the protein surface, colored from low (white) to high (red) contribution (kcal/mol). Right panels: Average energy of the protein-ligand interactions between PARP1 and different inhibitors. Only residues with an absolute energy contribution greater than 1 kcal/mol are shown. Energy components are broken down into van der Waals forces (vdW, blue), electrostatic interactions (El, purple), solvent polarization (PS, green), and non-polar solvent contribution (NPS, yellow).

[9] T. Karlberg, M. Hammarstrom, P. Schutz, L. Svensson, and H. Schuler, Crystal structure of the catalytic domain of human parp2 in complex with parp inhibitor abt-888, Biochemistry

**49**, 1056 (2010).

[10] A. R. Tejedor, A. Garaizar, J. Ramírez, and J. R. Espinosa, ‘rna modulation of transport properties and stability in phase-separated condensates, Biophysical Journal **120**, 5169 (2021).

[11] B. R. I. Miller, T. D. J. McGee, J. M. Swails, N. Homeyer, H. Gohlke, and A. E. Roitberg, Mmpbsa.py: An efficient program for end-state free energy calculations, Journal of Chemical Theory and Computation **8**, 3314 (2012).

[12] D. A. Case, H. M. Aktulga, K. Belfon, D. S. Cerutti, G. A. Cisneros, V. W. D. Cruzeiro, N. Forouzesh, T. J. Giese, A. W. Götz, H. Gohlke, S. Izadi, K. Kasavajhala, M. C. Kaymak, E. King, T. Kurtzman, T.-S. Lee, P. Li, J. Liu, T. Luchko, R. Luo, M. Manathunga, M. R. Machado, H. M. Nguyen, K. A. O’Hearn, A. V. Onufriev, F. Pan, S. Pantano, R. Qi, A. Rahnamoun, A. Risheh, S. Schott-Verdugo, A. Shajan, J. Swails, J. Wang, H. Wei, X. Wu, Y. Wu, S. Zhang, S. Zhao, Q. Zhu, T. E. I. Cheatham, D. R. Roe, A. Roitberg, C. Simmerling, D. M. York, M. C. Nagan, and K. M. J. Merz, Ambertools, Journal of Chemical Information and Modeling **63**, 6183 (2023).

[13] A. Onufriev, D. Bashford, and D. A. Case, Exploring protein native states and large-scale conformational changes with a modified generalized born model, Proteins: Structure, Function, and Bioinformatics **55**, 383 (2004), https://onlinelibrary.wiley.com/doi/pdf/10.1002/prot.20033.

[14] D. R. Roe and T. E. I. Cheatham, Ptraj and cpptraj: Software for processing and analysis of molecular dynamics trajectory data, Journal of Chemical Theory and Computation **9**, 3084 (2013).

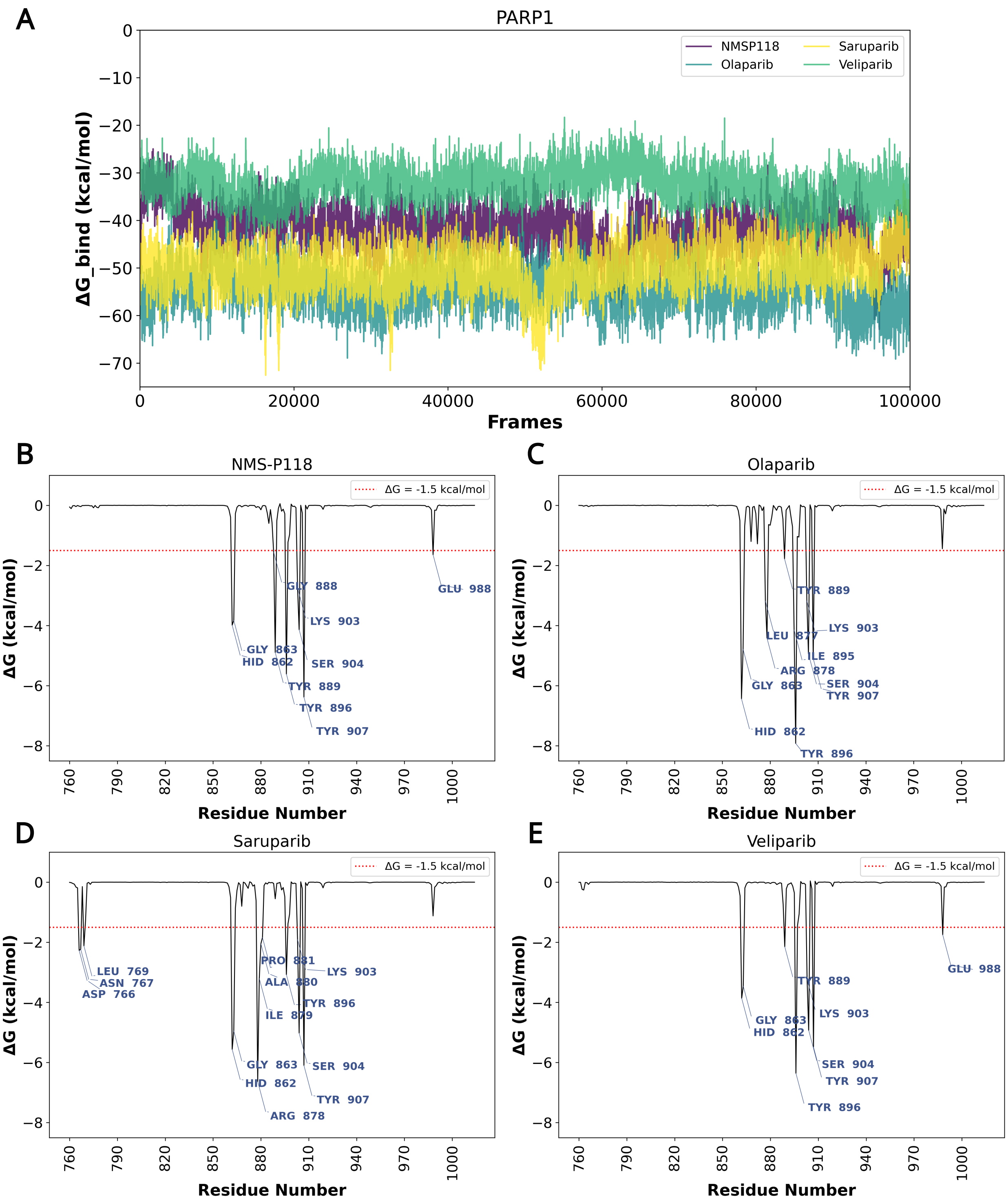


**FIG. S5:** (A) Binding free energy profiles calculated by MM-GBSA for NMS-P118, olaparib, saruparib, and veliparib in complex with PARP1 throughout the molecular dynamics trajectories. (B–E) Per-residue energy decomposition for NMS-P118 (B), olaparib (C), saruparib (D), and veliparib (E), highlighting the key residues that contribute most significantly to ligand binding. The red dashed line indicates the -1.5 kcal/mol threshold used to identify relevant interactions.

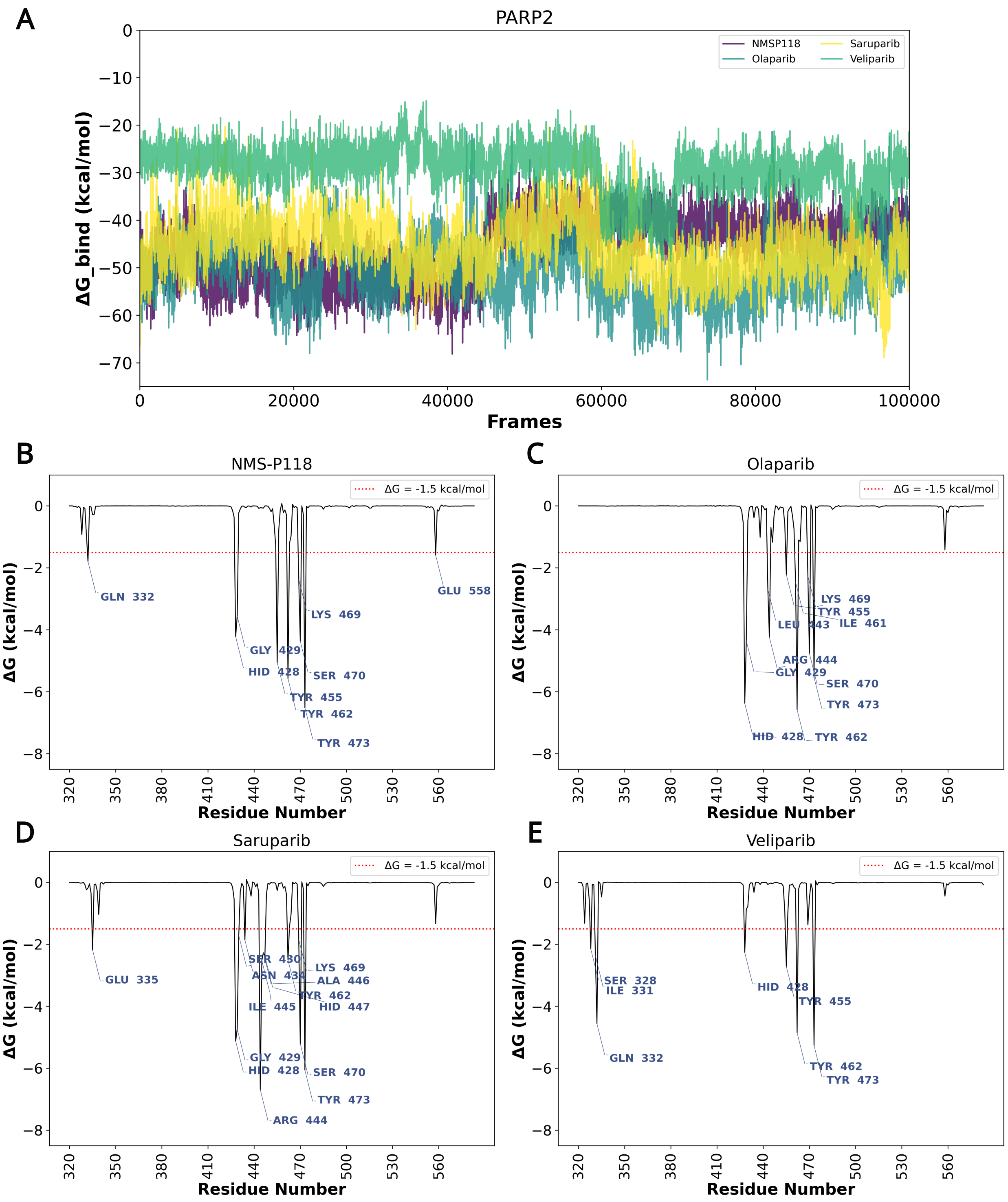


**FIG. S6:** (A) Binding free energy profiles calculated by MM-GBSA for NMS-P118, olaparib, saruparib, and veliparib in complex with PARP2 throughout the molecular dynamics trajectories. (B–E) Per-residue energy decomposition for NMS-P118 (B), olaparib (C), saruparib (D), and veliparib (E), highlighting the key residues that contribute most significantly to ligand binding. The red dashed line indicates the -1.5 kcal/mol threshold used to identify relevant interactions.

| | | PARP1 | | | | | | PARP2 | | | | | |
|---|---|---|---|---|---|---|---|---|---|---|---|---|---|
| **Ligand** | **AA** | **Residue** | **Donor** | **Acceptor** | **Distance** | **Angle** | **Frequency** | **Residue** | **Donor** | **Acceptor** | **Distance** | **Angle** | **Frequency** |
| **NMS-P118** | **SER** | 904 | O(P) | O(L) | 2,80 | 163,19 | 54% | 470 | O(P) | O(L) | 2,81 | 163,91 | 58% |
| | **GLY** | 863 | N(P) | O(L) | 2,89 | 162,62 | 54% | 429 | N(L) | O(P) | 2,87 | 155,37 | 55% |
| | | | - | - | - | - | - | | N(P) | O(L) | 2,89 | 156,47 | 50% |
| **Olaparib** | **GLY** | 863 | N(L) | O(P) | 2,83 | 162,82 | 86% | 429 | N(L) | O(P) | 2,86 | 161,50 | 73% |
| | | | N(P) | O(L) | 2,88 | 162,43 | 73% | | N(P) | O(L) | 2,86 | 155,01 | 71% |
| | **SER** | 904 | O(P) | O(L) | 2,80 | 163,46 | 70% | 470 | O(P) | O(L) | 2,80 | 165,87 | 71% |
| | TYR | 896 | N(P) | O(L) | 2,89 | 163,97 | 56% | - | - | - | - | - | - |
| **Saruparib** | **GLY** | 863 | N(P) | O(L) | 2,86 | 162,08 | 77% | 429 | N(L) | O(P) | 2,83 | 162,22 | 87% |
| | | | N(L) | O(P) | 2,86 | 161,51 | 63% | | N(P) | O(L) | 2,88 | 158,94 | 62% |
| | **SER** | 904 | O(P) | O(L) | 2,81 | 162,28 | 58% | 470 | O(P) | O(L) | 2,79 | 163,16 | 65% |
| | ILE | 879 | N(P) | O(L) | 2,85 | 151,26 | 58% | - | - | - | - | - | - |
| **Veliparib** | **SER** | 904 | O(P) | O(L) | 2,83 | 164,38 | 65% | - | - | - | - | - | - |

**FIG. S7:** Summary of hydrogen bonds detected between PARP1 or PARP2 and different inhibitors (NMS-P118, olaparib, saruparib, veliparib) in molecular dynamics simulations. Only interactions with an occupancy above 50% are shown.

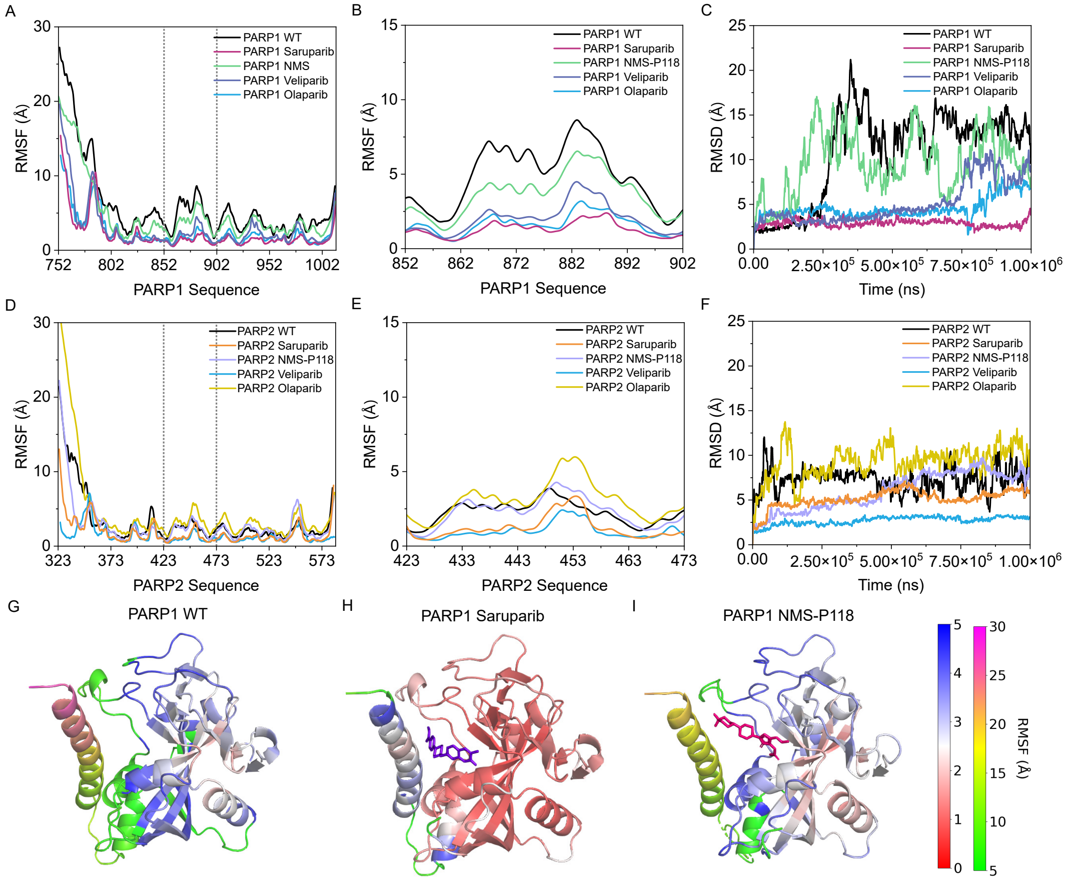


**FIG. S8:** Structural dynamics of PARP1 and PARP2 in the apo form and in complex with different inhibitors. Root Mean Square Fluctuation (RMSF) profiles of PARP1 for the wild-type (WT) protein and in complex with saruparib, NMS-P118, veliparib, and olaparib. (A) Global RMSF across residues 752–1002; (B) zoomed-in view of the flexible region spanning residues 852–902; (C) time evolution of the Root Mean Square Deviation (RMSD) for each system. Equivalent analyses for PARP2. (D) Global RMSF across residues 323–573; (E) zoomed-in view of residues 423–473; (F) RMSD over time for the different complexes and WT. Structural representations of PARP1 in the apo form (G), in complex with saruparib (H), and in complex with NMS-P118 (I). Proteins are shown in cartoon representation and colored according to per-residue RMSF values (see color bar), highlighting flexible regions and drug-induced stabilization effects.

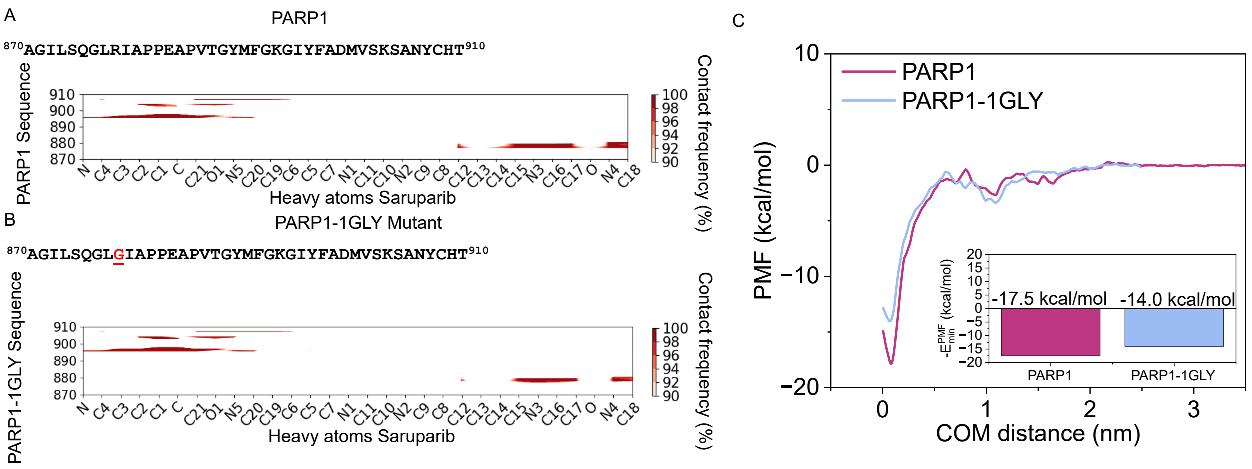

**FIG. S9:** Residue–ligand contact frequency maps between saruparib and PARP1 (A) and the 1GLY mutant (B). The PARP1 sequence is shown above each map, with the glycine substitution highlighted in red. Contact frequencies are calculated between heavy atoms of saruparib and zoom in the residues 870–910 of PARP1. The contacts are represented as contact frequency in percentage. (C) Potential of Mean Force (PMF) profiles for the unbinding of saruparib from PARP1 and the 1GLY mutant, computed as a function of the center-of-mass (COM) distance between the ligand and the protein. The inset shows the minimum binding free energy values derived from the PMF curves.

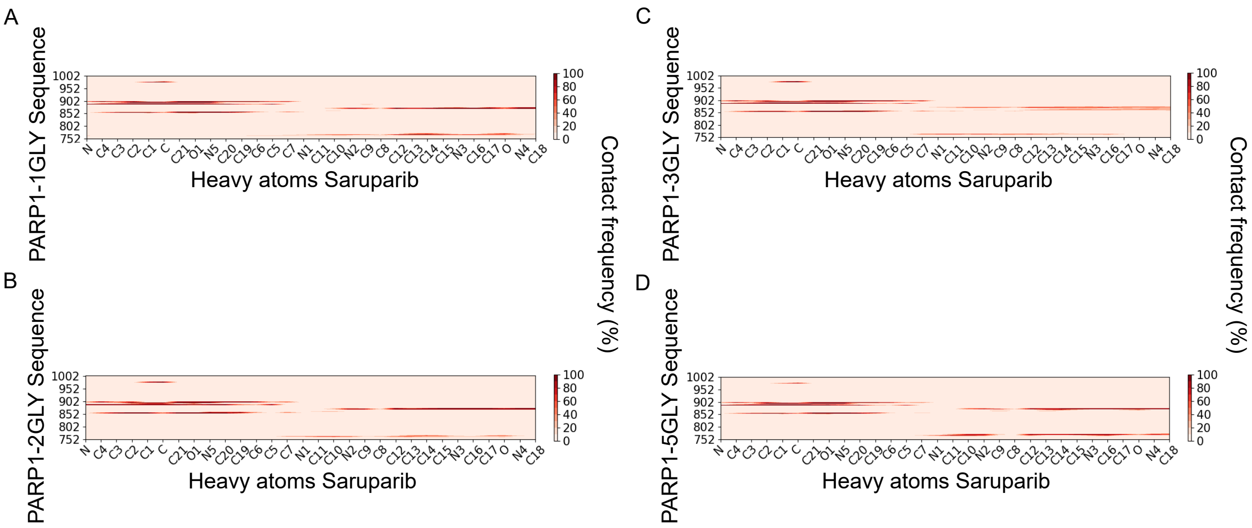

**FIG. S10:** Residue–ligand contact frequency maps between saruparib and 1GLY mutant (B), 2GLY mutant (B), 3GLY mutant (C) and 5GLY mutant (D). The contacts are represented as contact frequency in percentage.
